# TERribly Difficult: Searching for Telomerase RNAs in Saccharomycetes

**DOI:** 10.1101/323675

**Authors:** Maria Waldl, Bernhard C. Thiel, Roman Ochsenreiter, Alexander Holzenleiter, João Victor de Araujo Oliveira, Maria Emília M. T. Walter, Michael T. Wolfinger, Peter F. Stadler

## Abstract

The telomerase RNA in yeasts is large, usually > 1,000 nt, and contains functional elements that have been extensively studied experimentally in several disparate species. Nevertheless, they are very difficult to detect by homology-based methods and so far have escaped annotation in the majority of the genomes of Saccharomycotina. This is a consequence of sequences that evolve rapidly at nucleotide level, are subject to large variations in size, and are highly plastic with respect to their secondary structures. Here we report on a survey that was aimed at closing this gap in RNA annotation. Despite considerable efforts and the combination of a variety of different methods, it was only partially successful. While 27 new telomerase RNAs were identified, we had to restrict our efforts to the subgroup *Saccharomycetacea* because even this narrow subgroup was diverse enough to require different search models for different phylogenetic subgroups. More distant branches of the Saccharomycotina still remain without annotated telomerase RNA.

## 1. Introduction

The linear chromosomes of eukaryotes require a specialized mechanism for completing duplication. Most commonly this is achieved by a special reverse transcriptase, telomerase, that carries a specific RNA the template with telomeric sequence [1]. Most likely, this constitutes the ancestral state in eukaryotes. Despite its crucial function, telomerase has been lost several times in both animals (in particular insects) and possibly also in some plants [2]. In some cases, the ancestral telomere structure has been replaced by tandem arrays of DNA sequences that look much like heterochromatin and can be elongated by gene conversion. Specialized telomere-specific retrotransposons are at work in Drosophila [3].

The telomerase (holo)enzyme consists of two main components, a specialized reverse transcriptase (TERT) and a RNA component (TER) that provides the template sequence. In addition, there are usually multiple clade-specific accessory protein components [4]. Four conserved regions in TER, Fig. 1, are essential for telomerase activity: the template boundary element (TBE), the pseudoknot, and the template sequence itself are part of the the catalytic core. The fourth region, the trans activating domain, is involved in binding of TERT [5]. The three-way junction (TWJ) structure of this region is widely conserved at least between animal and fungal telomerase RNAs, where it is crucial for proper functioning [6]. The precisely defined template within TER is processively copied by TERT and regenerated, releasing a single-stranded DNA product [7].

**Figure 1.**
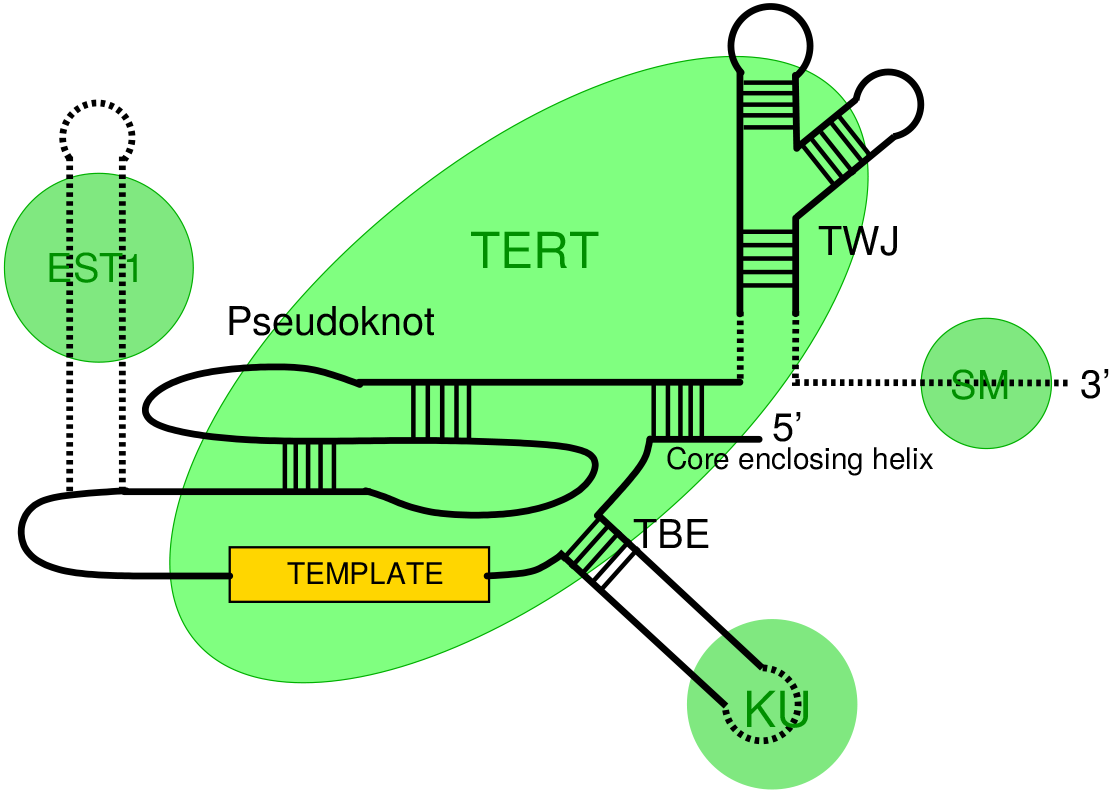
Schematic organization of TER. Contact regions for important binding sites are indicated by green circles (EST1, SM, KU). The green ellipse denotes the contact region with the reverse transcriptase (TERT). Other major features are the template, the pseudoknot region, the template boundary element (TBE) and the three-way junction (TWJ). Adapted from [8].

Telomerase RNA is highly divergent. The TER in ciliates [9], human [10], and budding yeast [11] have a length of about 150 nt, 438 nt, and ~1.3 kb, respectively. A TER more than 2kb in has been reported for *Candida glabrata* [12], which, interestingly, seems to lack a TWJ. TERs in other kingdoms of eukaryotes have been discovered only quite recently in plants [13, 14], excavates [15,16] and alveolates [17,18].

Despite their deeply conserved primary function and architectural similarities that seem to extend across eukaryotic kingdoms, TERs have turned out be very difficult to find by homology search even within phylogenetically relatively narrow groups. Within the animal kingdom, even surveys of vertebrates turned out to be non-trivial [19]. Echinoderm TERs were found by deep sequencing of *Strongylocentrotus purpuratus* RNA pulled down with the TERT protein [20] after homology based searches remained unsuccessful. This opened the door to identifying TERs from other sea urchins, brittle stars, and a crinoid [21]. Still, no TER from a protostome is known.

Within Fungi, the situation is similar: So far, TERs have been reported only for Ascomycota, while no candiates are known in Basidiomycota and any of the basal divisions. The TERs of Pezizomycotina and Taphrinomycotina share core features of vertebrate TERs. In particular, they have a fairly well-conserved secondary structure of the pseudoknot and the TWJ,and at least in these regions the sequence is sufficiently conserved for successful homology-based identification of TERs within these clades [22–24].The TERs known for Saccharomycetes, the relatives of budding yeast, on the other hand, are sometimes remarkably large and present little similarity in sequence and secondary structure to vertebrate or ciliate TERs.

To-date, yeast TERs have been reported for three phylogenetically narrow subgroups (*Saccharomyces spp.*[11,25], *Kluyveromyces spp.*[6,26,27], and *Candida spp.*[28,29]), as well as some individual species such as *Candida glabrata* [12] and *Hansenula polymorpha* [30]. These sequences are already too diverse for reliable sequence alignments. It is not surprising, therefore, that simple sequence-based homology searches have not been successful in identifying TER in the majority of the saccharomycete genome sequences to-date. Even protein binding sites that are functionally important in budding yeast [31] are not widely conserved. For instance, Ku or Sm binding sites seem to be absent in the TERs of filamentous fungi [4,22].

The obvious alternative is to increase the set of known TERs by finding homologs that are sufficiently similar to one of known yeast TERs, to allow the construction of multiple alignments of phylogenetically narrow subgroup. From these alignments, conserved elements can be extracted,which in turn form the basis for searches with tools such as fragrep [32] or infernal [33]. This strategy has been successful in previous searches for TER genes in both animals [19] and fungi [22], but so-far has not been successfully applied to Saccharomycetes.

Until very recently, a phylogenetically local approach to homology search was also hampered by the lack of a trustworthy phylogeny of the Saccharomycotina. Recent updates in the International Code of Nomenclature for algae, fungi and plants [34,35] have substantially restructured the classification of fungi in general and of Saccharomycotina in particular. With large-scale efforts to sequence fungal genomes underway, first phylogenomic studies provide a trustworthy backbone of Saccharomycotina phylogeny [36], which we largely confirmed with an independent analysis.

## 2. Materials and Methods

### 2.1. Phylogenomics of Ascomycotes

Annotated protein sequences for 72 yeast species were downloaded from RefSeq. Initially, `ProteinOrtho` [37,38] was used to identify an initial set of 21.289 ortholog groups. Only 193 of these contained representatives of all 72 species. We therefore included all 1666 ortholog groups that covered at least 67 species. We used `OMA` (2.2.1) [39,40] to decompose the `ProteinOrtho` groups further into clusters of 1-1 orthologs. This resulted in 6.295 groups of which 841 contained at least 67 species. This conservatively filtered data set was then processed with `Gblocks` [41] to remove uninformative and potentially error-prone parts of the alignment, resulting in a data set comprising 72 species and 248,581 characters. Phylogenetic trees were estimated with `RAxML` [42].

### 2.2. Ascomycote Telomerase RNAs

Telomerase RNA regions have been published for several *Saccharomyces* [11,25], *Kluyveromyces* [6,26,27], and *Candida* [12,28,29] species. Most of these published TER regions are collected in the telomerase database [43], which therefore provided a good starting point for our research. These sequences, however, are too diverse to construct multiple sequence alignments beyond the three genera individuall. This effectively prohibits the automated discovery of novel TERs beyond close relatives with the help of either blast [44] (using sequence information alone) or infernal (relying on a combination of sequence and secondary structure information).

Therefore, we explored different strategies to overcome the limitations imposed by the extremely poor sequence conservation of saccaromycete telomerase RNAs. The basic idea is to use common features of the TERs to extract candidates from the genomes that can be analyzed and then inspected further using different techniques.

First, we attempted to learn TER-specific sequence patterns using `MEME/GLAM2` [45], and also several machine learning techniques using *k*-mer distributions within sequence windows of the size of the known TERs. All attempts to learn from a training set covering the *Saccharomycetaceae* or all *Saccharomycotina* species failed.

There are several possible reasons. Machine learning methods crucially depend on a training and test sets, both positive and negative. In our case we have few positive samples, these have poorly defined features, and are very diverse as far as their sequences are concerned. It is unclear in this setting how a negative training set should be properly designed. The obvious choice of picking genomic sequence at random may be confounded by unintended strong signals, such as coding potential or repetitive sequence elements. It would appear that at the very least a more a careful construction of the positive and negative sets, and an appropriate normalization or scaling of the feature data will be required to make progress in this direction. Restricting the training phase to a more narrow phylogenetic range to reduce the inherent diversity of the training data, on the other hand, is infeasible due to the small number of known TER sequences.

The EDeN motif finder [46] was applied to 24 known TERs as positive set and 48 shuffled sequences as negative data. Only trivial sequence motifs such a poly-U stretch, presumably corresponding to part of the U-rich pseudoknot region, were found. Unsupervised clustering also remained unsuccessful.

### 2.3. Synteny-Based Homology Search

As an alternative strategy, we established a semi-automated workflow that aims at first extracting partially conserved RNA sequence-structure elements, which are then used to identify candidate loci. In response to the negative results of a direct pattern-based approach, we systematically used synteny to narrow down the search space in the initial phase. Starting from a whole genome alignment of phylogenetically related species, we used the positions of protein coding genes whose homologs are known to be adjacent in a closely related species to delimit the syntenic regions that are likely to contain a TER gene. These candidate regions were then analyzed in detail by means of pairwise or multiple sequence alignments. Whenever a global alignment of the entire candidate syntenic region did not yield a plausible alignment, we attempted to identify conserved motifs inside the syntenic region (usually the SM binding site and/or the template region, which is sometimes conserved between close relatives). Typically, these motifs were also sufficient to determine the correct reading direction of the TER candidate.

To identify known features in the candidate TER regions, we first constructed `infernal` [33] covariance models restricted to subgroups of Saccharomycetaceae covering only substructures, such as the Ku hairpin, Est1 binding site, and TWJ in the Saccharomyces and Kluyveromyces species. The alignments underlying the infernal models were constructed with the help of many software tools, including `locarna` [47], `mafft` [48], `mauve` [49], `MEME` [45] and `fragrep` [32], as well as manual curation. These models were then used for precise localization of conserved TER elements in species that were (a) taxonomically closely related, but not/only partially annotated in literature (*Saccharomyces uvarum*, *Saccharomyces sp. ‘boulardii’*, *Saccharomyces sp. M14*, *Saccharomyces eubayanus* or (b) phylogenetically located in the subtree spanned by the Saccharomyces and Kluyveromyces species (see Fig. 2). Both the `ViennaNGS` [50] suite and custom Perl/Python scripts were used for handling and conversion of genomic annotation data.

**Figure 2.**
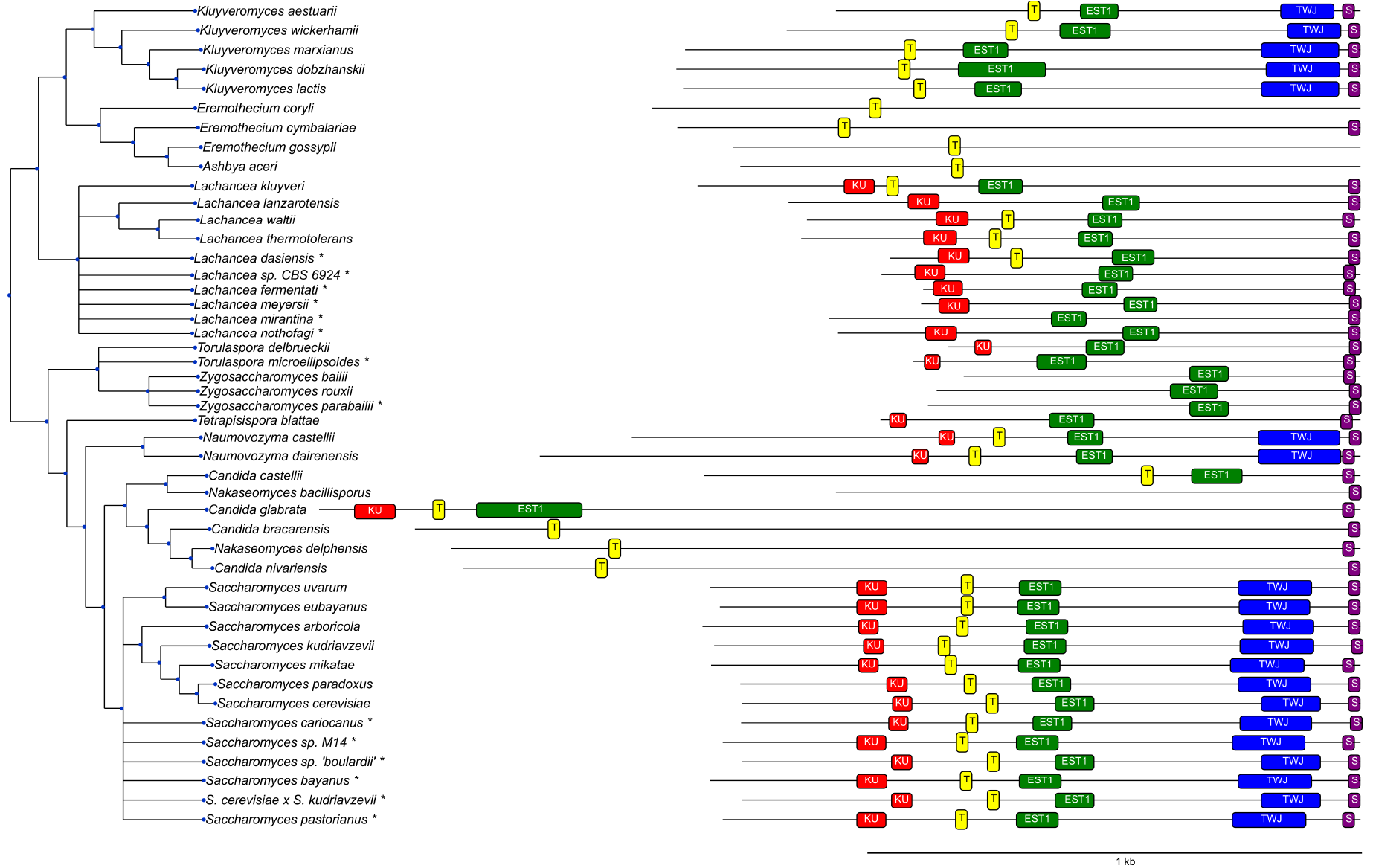
Features identified in TER sequences. **KU)** ku binding hairpin, **T)** template region, **EST1)** Est1 binding site, **TWJ)** three-way junction, **SM1)** SM1 binding site. Elements not shown are either not present in the corresponding species (e.g. the TWJ in *C. glabrata*) or could not be located with reasonable certainty. Species marked by * are not part of the phylogenetic tree and were placed next to their closest related neihbour based on the similarity of their TER sequences.

We then extracted a sequence corresponding to the most closely related TER sequence as initial estimate of the full-length TER gene. We used `mafft` [48] to produce initial sequence-based alignments of candidate regions, which were then realigned with `locarna` [47] to obtain RNA structural alignments. The latter was used with its free-end-gaps option, in particular in those cases where `mafft` was not sensitive enough to reliably estimate the TER boundaries. Conversely, `mafft` was able to identify and correctly align highly conserved subsequences, providing reliable anchors for the more divergent sequence regions. While `locarna` is good at finding locally conserved structures in the whole alignment, we expected only parts of the TER sequences to be structurally conserved. Typically multiple iterations of refinement of the TER boundaries were required to obtain the final TER candidate sequence.

Following this approach, we could localize TER regions for several members of the *Saccharomycetacea* clade. Subsequent alignment of candidate regions with known TERs allowed for exact localization of TERs.

### 2.4. Search for Candidates Using Telomere Template Sequences

The scope of the synteny-based approach is limited because fungal genomes are subject to frequent genome rearrangements at the time-scales of interest. We therefore attempted to identify candidate regions containing the template sequence for the telomere repeats. (See [51] for a comprehensive review of the characteristics of different telomeric repeats.) In genomes for which these sequences have not been reported, we searched chromosome ends for telomeric repeats. Unfortunately, most genome assemblies are not on chromosome level or do not include the telomere regions, hence we only succeeded to newly identify the template region of *Ashbya aceri* and *Eremothecium cymbalariae* this way. For the latter species, the pertinent information is available in [52], although the telomeric repeat is not explicitly reported. In addition, we used the published telomere sequences from the telomerase database [43].

We used the concatenation of two copies of telomeric repeat sequence as query for a `blast` [44] search against the whole genome (in case of longer, complex repeats) or against the syntenic region for shorter repeats. Other template regions were identified with by aligning them to known sequences and/or `blast` searches of known template regions in closely related species. A typical feature of the template region, which helped us to verify our hits, is the fact that it usually contains a few nucleotides repeated at both the beginning and the end of the template region [12].

### 2.5. Blast Pipeline

`blast` [44] is by far the most commonly used tool for homology search. While it has been reported to have limited sensitivity for telomerase RNAs in previous studies [19,20,32], it has contributed significantly to the identification of the TER sequences in other ascomycete clades [22,24]. Here we used a set of known TER regions as `blast` queries that comprises all Saccharomycetales TER regions that we found in literature, as well as all TERs newly identified in the contribution. As targets for `blastn` (with default parameters) we used the full genomes of species that are featured at the `NCBI refseq` database within the Saccharomycetales group (Taxonomy ID: 4892). The resulting blast hits were then filtered for E-values (*E* < 0.1), a minimum alignment length of 25nt and a minimum identity172of 60%. In addition, all hits on known telomeric regions were excluded. From the hits in genomes with known TERs we computed the empirical false positive rate and found that the alignment length proved to be the most informative parameter. It has therefore been used to evaluate the reliability of hits, given their score.

The `blast` pipeline also contributed to the identification of the TER boundaries in some of the unannotated genomes. In cases were we initially chose the boundaries of our queries too generously and included neighboring coding regions or regulatory elements, the blast pipeline returned “false positive” hits. Thus, whenever multiple false positive hits in the beginning or the end of the query sequence occurred, we rechecked and, if necessary, improved the boundaries of the TER region.

## 3. Results

### 3.1. Phylogenomics of Saccharomycotina

The phylogenetic trees obtained of our phylogenomic analysis of the Saccharomycetales is essentially congruent with the one reported by Shen *et al.*[36], see the Appendix for more details. For consistency, we adopted the phylogenetic tree published by Shen *et al.*[36] as the basis for presenting our results.

### 3.2. Survey of TER Genes in Saccharomycotina

We initially screened 52 ascomycote genomes. Predominantly sequence-based methods (`blast`, but also `meme, glam2`, and `infernal`) only contributed TERs from close relatives of baker’s yeast. The `blast` pipeline was applied to all NCBI genomes Saccharomycetales, the subclade containing all known Saccharomycotina genomes. With the exception of the TER in *Ogataea parapolymorpha*, a very close relative of the known *Ogataea polymorpha* TER [30] all new sequences we found within the Saccharomycetaceae. We therefore restricted a more detailed analysis to this clade.

We found credible TER sequences in 46 of the 53 Saccharomycetaceae. Most of these TER sequences could be detected only after a short candidate region had been identified based on synteny. To our knowledge, at least 27 of these have not been reported previously.

### 3.3. Features of TER in Saccharomycetacea

In order to better understand the TER and its evolutionary constraints at least within the Saccharomycetacea we performed a detailed analysis of their structural features. Table 1 summarizes the results of the homology search and the functional features of the candidate TER genes. A graphical overview is given in Fig. 2.

**Table 1.**
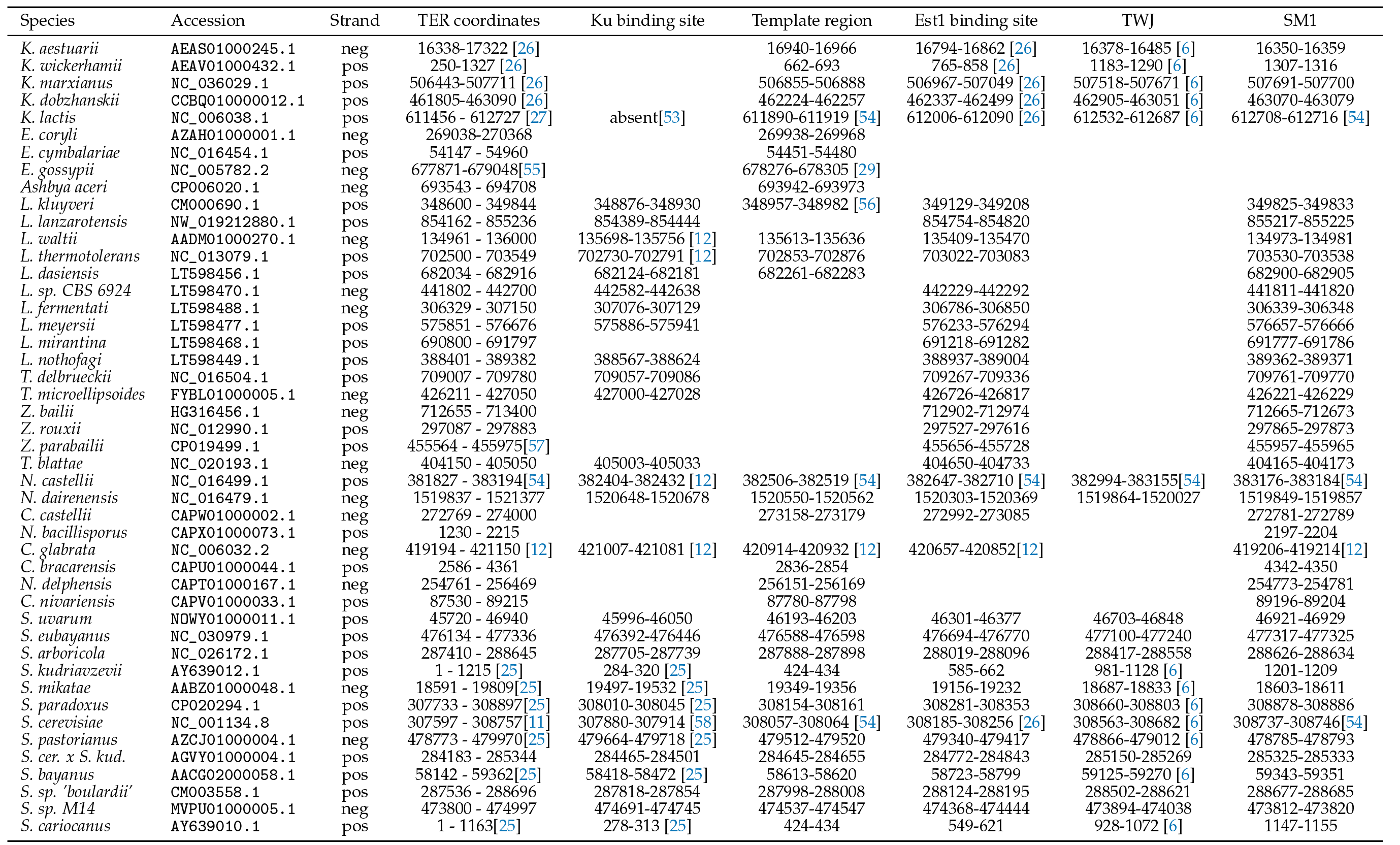
Overview of conserved telomer substructures in Saccharomycetacea, as identified by the combined synteny/covariance model pipeline. The 3’ end is defined as 10 nt downstream of the SM binding site. The 5’ end is approximate. Citations refer to publication in which the sequence and/or the coordinates of respective features are reported explicitly. These annotations form the basis of Figure 2.

The exact genomic positions marking the 3’ and 5’ ends of the TER RNA are difficult to determine without additional experimental evidence. The 5’ ends are therefore approximate. The 3’ end of the mature TER is produced by splicing in most Ascomycota [24,59,60]. This mechanism, however, was lost at some point during the evolution of the Saccharomycotina. It has been reported in the Candida group and for *Ogataea angusta* (previously *Hansenula polymorpha*), but it is missing in Saccharomyces and Kluyveromyces [24]; hence we expect that the splicing-based 3’-end processing was lost prior to the divergence of Saccharomycetacea. Indeed, no indication of a splice site was found for any of the TER sequences included in Table 1. We therefore used a position 10 nt downstream of the SM binding motif as approximation of the 3’ end in Table 1.

Several of the features listed in Table 1 have been discussed in some detail in the literature. Not all of them were found in all the candidates reported here. This may, in some cases, be explained by sequences that are too divergent to be detected. In other cases, most likely the function is not preserved. Unfortunately, many studies report neither complete sequences nor coordinates, making it effectively impossible to accurately compare their results with the annotation reported here. References are included in Table 1 if sufficient information was included to locate the features unambiguously.

No Ku binding hairpin was recovered in *Kluyveromyces* or the *Eremothecium* species. This is not unexpected since there is experimental evidence that neither the Ku binding hairpin nor its function is present in *K. lactis* [53]. The putative Ku binding hairpin reported for *Candida glabrata* in [12] lacks experimental support and contains long insertions that made it impossible to include it in our covariance model. Furthermore, this region of the TER sequence is very poorly conserved in the closest relatives of *C. glabrata*. While the TER of *C. glabrata* is among the longest known members of this gene family [12], its close relative *C. castellii* features a TER that has been shortened drastically in its 3’ half, with only ~200 nt separating the EST1 and SM1 binding sites. Furthermore, the sequence GCUA, which is conserved in most known Ku binding sites, is not present within 600nts upstream of the template region. The most likely explanation is that the TER of *Candida castellii* (which like *Candida glabrata* does not belong to the monophylogenetic genus *Candida*, see Appendix) does not bind Ku. Of course, we cannot rule out without further experimental data that the motif has diverged beyond our ability to recognize it.

In a few species we failed to identify the template region. In these cases (*Lachancea*, *Zygosaccharomyces* and *Torulaspora* species and *Nakaseomyces bacillisporus*) the telomeric repeat sequence is not known and seems to be very different from both the fungal consensus sequence `TTAGGG` [22] and the telomeric sequences found in closely related species.

The EST1 binding site could not be identified in *Eremothecium* species, *Lachancea dasiensis* and in the *Candida glabrata* group, even though it has been published for *Candida glabrata*. While an EST1 binding site is present even in the more distantly related genus *Candida* [29], this motif is intrisically too variable to be unambiguously recognizable in distant relatives. This pertains to both its sequence and the its base-pairing patterns.

Consistent with [12], we found no plausible secondary structure for the TWJ in *C. glabrata*, although the respective region of the sequence contains the highly conserved sequence `AATA`. It is worth noting in this context that the telomerase of the ciliate *Tetrahymena* has a stem-loop structure in place of the threeway junction [61]. TERs of the *C. glabrata* group thus may also have a functional trans-activation domain, albeit with an aberrant structure. Our TWJ covariance model, which was constructed from *Kluyveromyces* and *Saccharomyces* sequences only, also failed to detect a TWJ in *Eremothecium* and *Lachancea*. It remains an open question whether TERs of these species have a TWJ with a diverged structure that is just beyond our ability to detect, or whether trans-activation is achieved by different means.

The sequence of the SM binding motif AATTTTTGG is perfectly conserved throughout much of the Saccharomycetaceae, with the notable exception of *K.lactis* [54] and additional small variations in other *Kluyoveromyces* species, see Fig. 3. We could not find this motif in species of the genus *Eremothecium*and the highly related species *Ashba acerii*.

**Figure 3.**
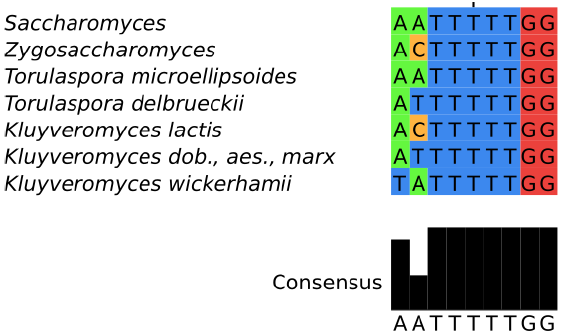
Alignment of the core SM-binding site motif. The common pattern of most Saccharomycetaceae is shown on top, species-specific variants are listed below.

## 4. Discussion

Although we succeeded in detecting 27 previously unknown TER sequences in Saccharomycetaceae, the main take-home message is of this contribution is that homology search can be a terribly difficult problem. Although yeast TERs are quite long and fulfil a well-conserved function, their sequences are very poorly conserved. In this respect, yeast TER behaves much like the majority of long non-coding RNAs, which are also poorly conserved in sequence but often are evolutionary quite well conserved as functional entitied, see [62] for a recent review.

The “`blast graph`” in Figure 4 highlights the practical problem. Sequence comparison methods identify homology only in closely related species. A comparison of Figure 4 and a corresponding graph based on the previously published TER sequences only (see Online Supplemental Material) shows that the larger set of queries identifies many additional connections and thus improves the situation at least within the Saccharomycetacea. Even within the clade, however, we have been unable to confirm the candidate hits in *Kasachstania*. The tree in Figure A1 indicates longer branch lengths leading to *Kasachstania*; it appears that the accelerated evolution of these genomes is already sufficient to hide the TER genes from our homology search methods.

**Figure 4.**
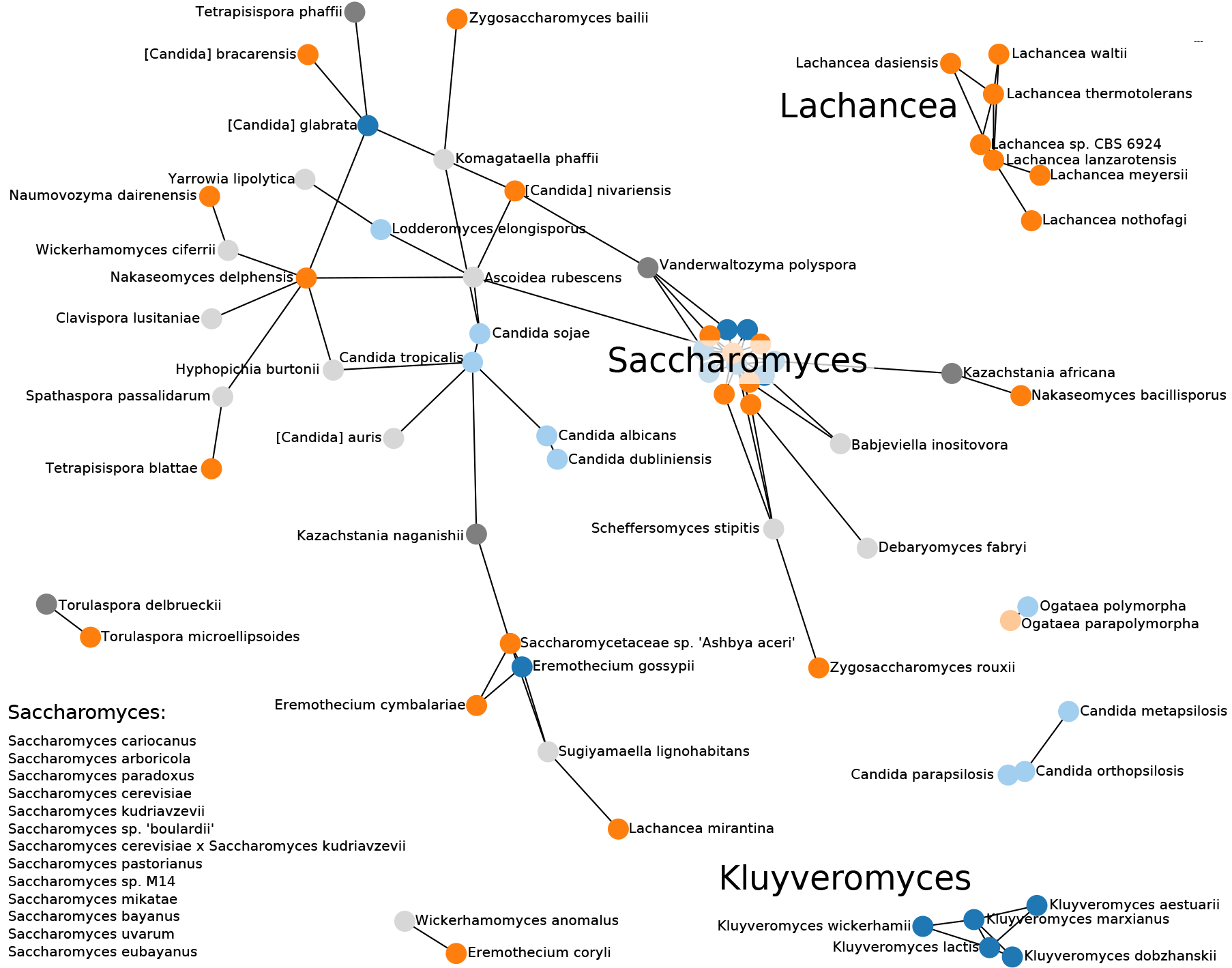
Summary of the blast-based survey of TER genes. Blue nodes show TERs described in literature, orange nodes represent TERs that we identified, and grey nodes are additional candidates for which we could not validate characteristic features. TERs outside the Saccharomycetaceae group are presented in light colors. The length of the edges are weighted by the inverse of the length of the `blast` hit. Note that distances in drawing between nodes not connected by an edge are not indicative of their evolutionary distance.

**Figure A1.**
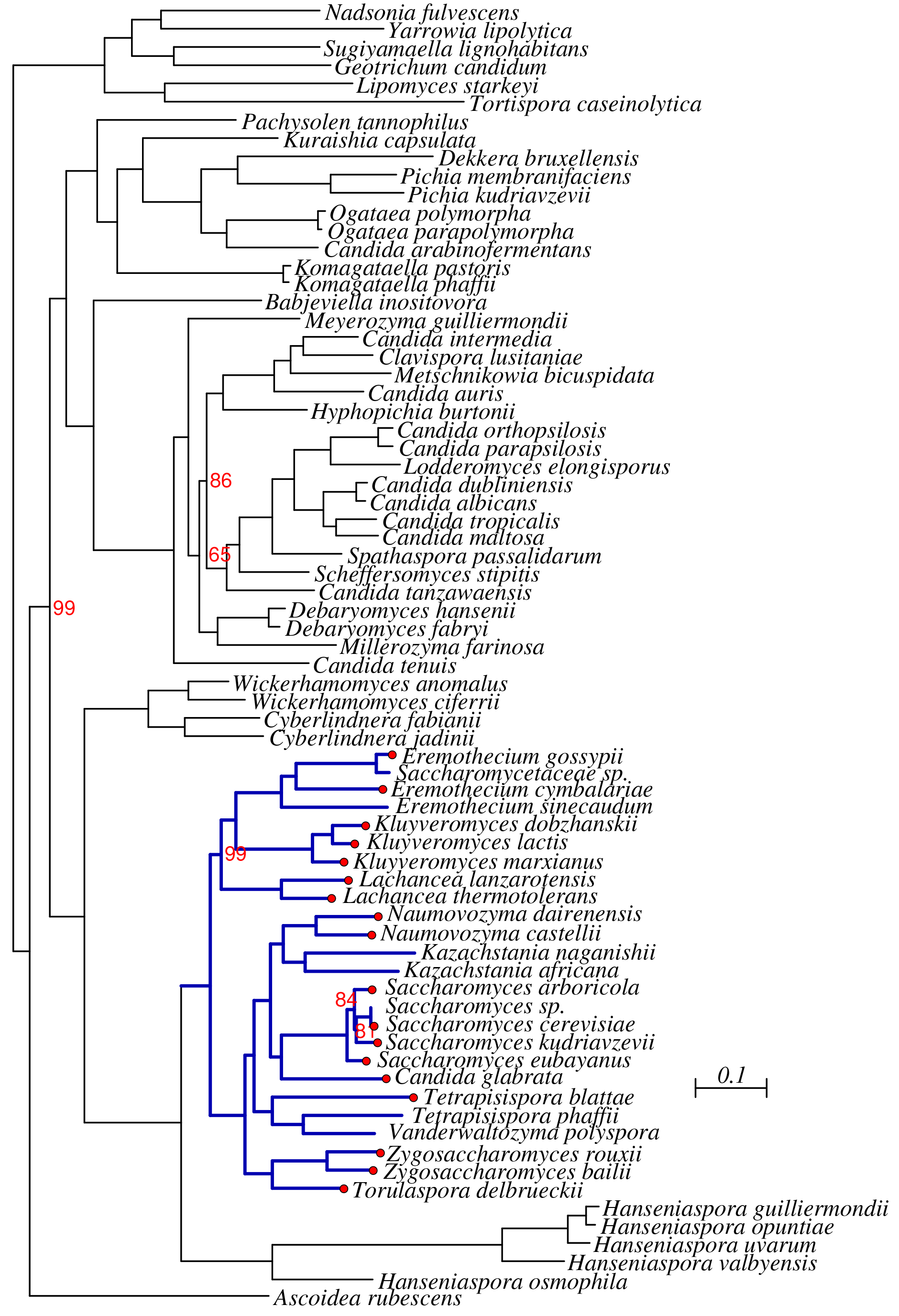
Phylogeny of the Saccharomycetales. Bootstrap support is 100% unless otherwise indicated. The Saccharomycetacea are indicted in dark blue. A red dot at tip of the tree indicates a TER sequences listed in Table 1.

While the direct sequence-based search against complete genomes was not very successful, we observed that the synteny-based approach worked remarkably well. This is not entirely unexpected, since the restriction to the interval between a pair of coding genes effectively reduces the size of the target from several million nucleotides to a few thousand. Unfortunately, the applicability of synteny-based methods is limited to relatively narrow phylogenetic scales. On longer time-scales, genome rearrangments are likely to disrupt syntenic conservation. A systematic exploitation of synteny similar to the work described here for Saccharomycetacea would most likely be successful in a survey for TER in the Candida group. In fact synteny has been employed to find some of the known TERs in this clade [29].

The study presented here was largely conducted using publicly available tools complemented by some custom scripting. It also highlights the need for customized tools to conduct difficult homology searches. In particular, specific alignment tools and viewers to efficiently evaluate the synteny-based candidates relative to known template sequences and alignments of the better conserved regions would facilitate the manual curation efforts, which we found to be indispensible.

Finally, it remains on open question whether direct machine learning methods can be adapted as homology search tools, and if so, whether such a strategy can be more effective than sequence comparison methods. It is likely that such efforts failed so far because of the difficulties inherent in the construction of a suitable negative training set that is not confounded by frequent genomic features such as coding sequence. Furthermore, the small number of positive samples was presumably insufficient to capture the full variability of TER sequences.

Complementarily, a phylogenetically dense sample of TERs that are sufficiently similar to support global sequence aligments might help to better understand the rapid divergence of TER sequences. This may be helpful not only to identify informative features for machine learning applications, but may also help to design modified sequence comparison algorithms that better reflect the peculiarities of rapidly evolving long non-codign RNAs. In this contribution we have provided such a set of TERs for the Saccharomycetaceae.

## Supplementary Materials

Machine readable Supplemental Information, in particular accession numbers, TER sequences, alignments of conserved features, and covariance models are available at http://www.bioinf.uni-leipzig.de/Publications/SUPPLEMENTS/18-048/.

## Acknowledgments

The research reported here is the outcome of a PhD level seminar taught by PFS at Univ. Vienna in the fall term 2017/18.It was funded in part by the Doktoratskolleg RNA Biology at Univ. Vienna, the Austrian Science Fund (SFB 4305-B09, I 2874-N28, I-1303 B21), Sinergia (CRSII3_154471/1), and the German Federal Ministery for Education and Research (BMBF 031A538A, de.NBI/RBC).

## Author Contributions

PFS and MTW conceived the study. MW, BT, RO, and MTW analyzed the TER sequences and structures, AH conducted the phylogenomic analysis, JVdAO and MEMTW contributed machine learning approaches. All authors contributed to writing the paper and approved of the submitted version.

## Conflicts of Interest

The authors declare no conflict of interest.

## Appendix A. Phylogenomics of the Saccharomycetales

The maximum likelihood tree obtained from 841 orthologous groups of proteins present in at least 67 of the 72 species is shown in Fig. A1.The phylogeny is nearly identical to the tree reported in [36]. In particular, it provides strong support for monophyletic Saccharomycetacea (comprising in particular the genera *Saccharomyces* and *Kluyveromyces*), and the Candida group. Noteworthy, “*Candida glabrata*” is nested within the Saccharomycetacea as a close relative of *Saccharomyces* rather than appearing as member of the Candida clade.

